# Resolving Host-Episymbiont Interaction Dynamics through Continuous Cultivation

**DOI:** 10.64898/2026.05.01.722272

**Authors:** Alex Grossman, Jacey Weng, Adam D. Silverman, Batbileg Bor

## Abstract

Patescibacteria are an elusive linage of “microbial dark matter” bacteria predicted to represent ∼25% of total bacterial diversity. Despite this abundance and ubiquity, these organisms are challenging to cultivate, resulting from their specialized episymbiotic lifestyle. All cultivated representatives to date, predominantly composed of Saccharibacteria from the oral microbiome, depend on cognate prokaryotic hosts for growth and reproduction. Studying the growth dynamics of episymbiotic bacteria and their hosts in batch cultures has suggested that many episymbionts initially reduce host populations, and that hosts eventually adapt to episymbiont stress after serial passaging. However, discontinuous batch cultures do not reflect natural interactions between these organisms due to their drastically different growth rates. An episymbiont requires several (∼2-4) serial passages alongside its host to reach the high cell densities needed to impact host growth, which complicates investigation of host inhibition and adaptation to episymbiont stress. To describe these dynamics accurately, we utilized continuous culture via small-scale Raspberry Pi powered bioreactors, called Pioreactors. Within a bioreactor, host bacteria can be cultivated at a consistent growth rate indefinitely, providing the perfect substrate for cultivation of model Saccharibacteria. Quantification of time until host crash, crash severity, time until recovery, and stable co-culture density provides mechanistic ways to describe episymbiont-host interactions. First, we used these techniques to compare episymbiont infection by three different episymbionts, revealing distinct infection patterns ranging from mild inhibition with rapid host adaptation, to rapid host collapse followed by “arms race” oscillation dynamics. Then, bioreactors were used to quantify the episymbiotic role played by a known host-binding type 4 pili (T4P-2), demonstrating that loss of long-distance host binding significantly delayed the host crash without altering general crash dynamics. These experiments reveal that episymbionts can have drastically different effects on bacterial communities and provide the tools necessary to describe strain/species differences and molecular interactions.

**Importance:** Episymbiotic Patescibacteria represent one of the largest branches of life on Earth, as well as one of the least understood. Furthermore, because Patescibacteria can manipulate their hosts growth and morphology they have immense ecological potential to be shaping the communities they occupy, both environmental and microbiome-associated. Our study highlights for the first time the potential of small-scale continuous cultivation for studying episymbiotic interactions that cannot be captured in discontinuous cultures. Herein we used these techniques to interrogate inter-species variation in host inhibition potential and to determine how loss of a long-distance episymbiosis factor mechanistically alters the cycle of episymbiont infection; however, this cultivation platform will enable researchers to answer many new questions about these ubiquitous host-episymbiont interactions.

## Introduction

Episymbiotic Patescibacteria are elusive, ultrasmall microbes from an ancient branch of the bacterial family tree that have seemingly become an adaptive radiation specializing in obligate episymbiosis (1). They possess streamlined genomes and grow in physical contact with the surface of larger bacteria or archaea (2). Very little is known about the ecological processes performed by these microbes (3). While only recently discovered to occur in bacteria, episymbiosis is not a rare microbial lifestyle. Patescibacteria represent a large portion of the “uncultivatable” microbial dark matter and ∼25% of total bacterial diversity (4, 5). Within the last decade, several host-episymbiont co-cultures, predominantly Saccharibacteria from the human oral microbiome, have been successfully cultivated and used to demonstrate the profound ways episymbionts impact host microbes’ growth and morphology (6, 7). These studies suggest that episymbionts exert powerful effects on microbiome function and composition, worthy of understanding. However, many bacteriological techniques are complicated by episymbionts’ host requirements. Growing two organisms with different reproductive tempos in a closed system prevents study of long-term population dynamics. Episymbionts grow significantly slower than their hosts, and episymbiont growth halts when the host reaches stationary phase (8, 9). Repeated subculturing of co-cultures, or passaging, can enrich episymbionts enough to detect host growth inhibition (7), and passaging beyond that point often results in episymbiont-resistant host adaptation (8). To understand the role of Saccharibacteria within the human microbiome, we must understand the dynamics of host decline and adaptation; however, serial passaging makes these phenotypes intractable. Continuous cultivation in a carefully controlled bioreactor system with a steady flow of nutrients provides a more natural environment by enabling both populations to grow continuously (Figure 1A) (10). This kind of bioreactor is referred to as a chemostat because it maintains a steady chemical composition via constant culture dilution. Furthermore, the recent development of small bioreactors, like the Raspberry Pi powered Pioreactors used in this study (11, 12), reduces experimental costs and enables incorporation of chemostats into existing anaerobic/microaerophilic cultivation chambers capable of Saccharibacteria cultivation (Figure 1B).

**Figure 1.**
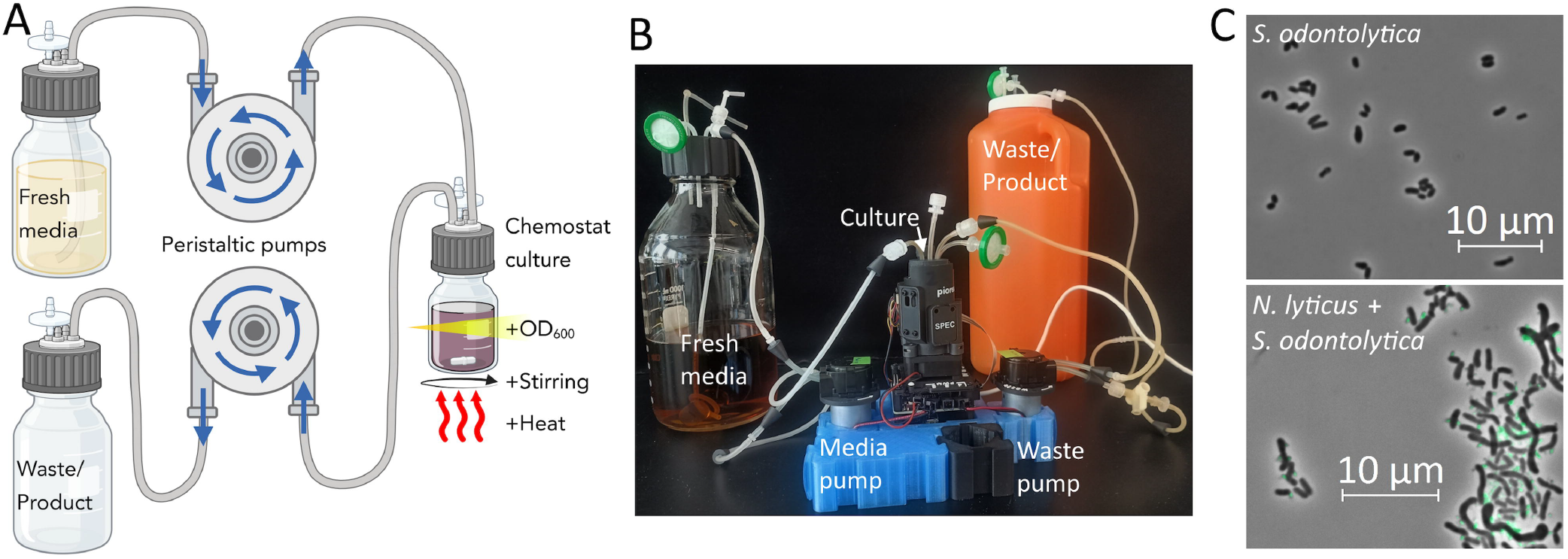
Cultivation of episymbiont co-cultures within a bioreactor. **(A)** A bioreactor diagram depicting the components required and the direction of flow. **(B)** Our experiments specifically adapted small Pioreactor platforms because of their size and modularity. These systems house the culture flask and sensors inside the central black tower which is flanked by the peristaltic pumps that power media addition and culture removal. **(C)** Our model host bacterium, *S. odontolytica*, is native to the oral microbiome and typically grows as pleomorphic rods. When grown in co-culture with *N. lyticus* (expressing green fluorescent protein for easy visualization), the episymbionts attach themselves to *S. odontolytica* surfaces and induce host elongation and clumping.

To probe these episymbiotic interactions, our experiments utilized a host organism isolated from the human salivary microbiome, the Actinomycete *Schaalia odontolytica* strain XH001 (Figure 1C) (13). *Actinomyces*/*Schaalia* complex microbes are almost universally present commensals in the oral microbiome, occur at high abundance, (14) and are often acquired at an early age (15, 16); however, they can also act as opportunistic pathogens contributing to Actinomycosis, blood infections, and polymicrobial infections (16, 17). Furthermore, studies of persistently inflamed tissues (periodontitis, gingivitis, inflammatory bowel disease) show declines in *Actinomyces*/*Schaalia* populations and increases in the Saccharibacteria that are predicted to use them as hosts, highlighting the complexity of these polymicrobial infection communities (18–22). *S. odontolytica* XH001 uniquely can host, or be infected by, three different cultivatable Saccharibacterium species from the family *Nanosynbacteraceae*; *Nanosynbacter lyticus* (HMT-952) strain TM7x, HMT-352 strain TM7-008 (ANI_*N*. *lyticus*_ 83.8%), and HMT-488 strain BB004 (ANI_*N*. *lyticus*_ 92.8%) (6, 8, 23). While all three episymbiont species share a common host organism and are isolated from the oral microbiome, previous research has shown that they induce different morphologies in host cells and differ in their competitive fitness (23). Specifically, TM7-008 is a strong competitor, BB004 is a weak competitor, and *N. lyticus* is intermediate. Additionally, to provide a reductionist example of how chemostat cultivation enables study of episymbiosis, we used a previously identified *N. lyticus*Δ*pilA2* mutant that is unable to form a host-binding type IV pili (23). Continuous cultivation enables analysis of how each episymbiont’s host infection cycle differs and how those differences impact host populations.

## Results

To interpret the cycle of episymbiont infection and host adaptation in continuous culture, new metrics had to be developed. First, host cells are grown until they reach a stable density and growth rate, resulting in a **host steady-state** (Figure 2A). The host steady-state represents the point when host cell monocultures growth perfectly matches the dilution factor of the chemostat. In our experiments, 15 mL cultures continuously fluxed media at a flow rate of 4.56 mL/hour, resulting in a dilution factor of 0.30 per hour. To visualize the impact of episymbionts on host growth after infection, the host steady-state was used to normalize all later OD_600_ readings (Supplemental Figure 1A-B). Next, isolated episymbionts (∼4*10^7^ cells) were injected into the bioreactor at roughly a multiplicity of infection (MOI) of 0.01 episymbionts/host. For our purposes, episymbionts are assumed to have no impact on optical density because their light scattering is negligible relative to host cells (9). After infection, densities either remained constant or showed slightly enhanced growth for a number of hours that we termed “**time until crash**”. Eventually, episymbionts reach sufficient density to begin inhibiting host growth, causing the optical density to start a precipitous, logarithmic decrease that we termed the **crash phase**. Two metrics were used to quantify crash phase severity: **crash rate** was determined by fitting a linear function to the period of peak decline, and **surviving population** was estimated using the lowest density achieved before **population recovery** began. Eventually, host adaptation and dilution of episymbionts allows host populations to begin increasing, resulting in a **recovery phase**. The time between crash and host recovery was termed “**time until recovery**” and used as a proxy for rate of host adaptation. After the recovery phase, host-episymbiont co-cultures eventually reach a new steady-state termed **stable-symbiosis density**, at a lower OD_600_ than the host steady-state (Figure 1A).

**Figure 2.**
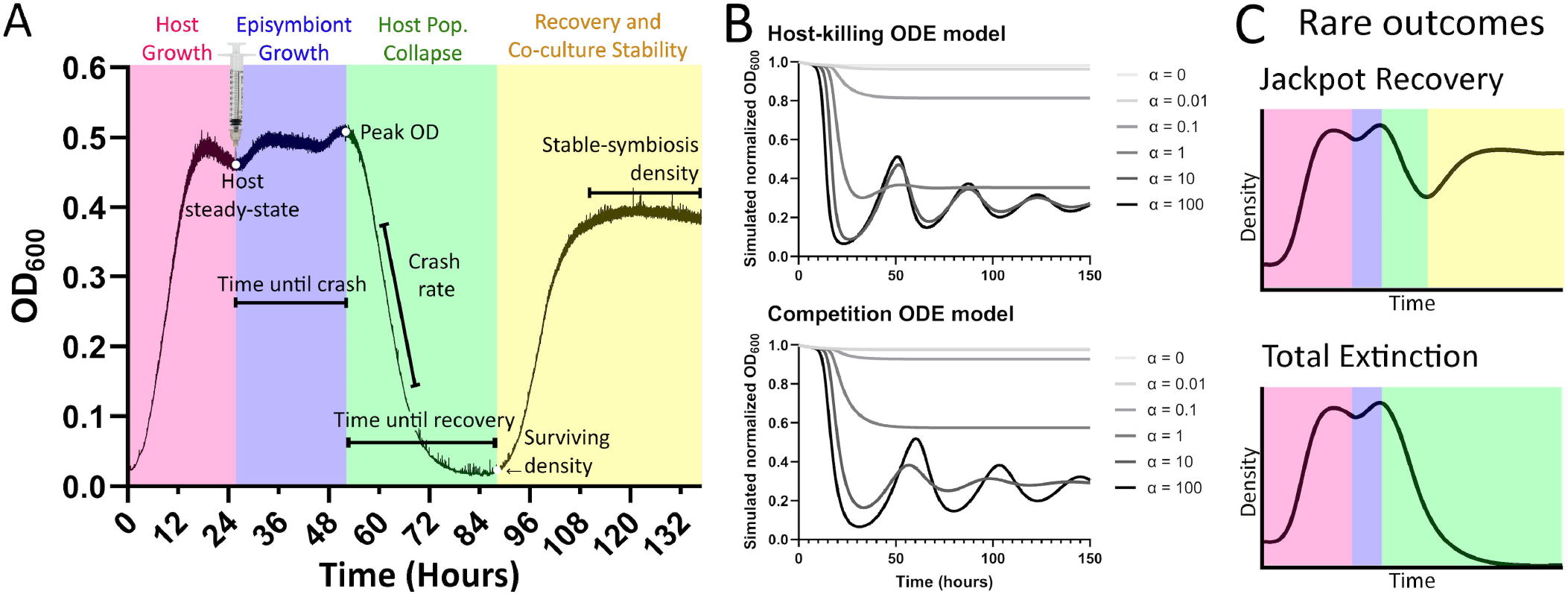
Model cycle of episymbiotic infection by a Saccharibacteria. **(A)** Schematic diagram of the standard episymbiont infection cycle as observed within a chemostat, divided into 4 phases; host monoculture growth until stability (pink), episymbiont growth on continuously growing host (blue), the collapse of the host population from episymbiont stress (green), and finally host adaptation/recovery (yellow). Important metrics such as OD_600_ values at phase transitions, time until crash, time until recovery, and crash rate are highlighted. **(B)** Two different ordinary differential equation (ODE) models were designed to simulate the observed cycles of infection; a host-killing model (top) that assumes episymbionts kill associated hosts at rate α and a competition-based model (bottom) that assumes episymbionts inhibit host growth at rate α. **(C)** Over the course of our experiments, two “rare outcomes” outside the standard models occurred. A jackpot recovery occurs when the host population adapts to become resistant before drastic growth inhibition, and a total extinction occurs when host growth is inhibited so strongly that all host cells are diluted out of the bioreactor or killed.

In parallel with our experiments, we developed two ordinary differential equation (ODE) models to qualitatively reproduce the host-episymbiont growth dynamics observed in our experiments (see Supplemental Files 1 and 2 for model details). Our first model assumes that the episymbiont actively kills the host (with second-order rate constant α) and that episymbiont growth rate is proportional to host population, like a Lotka-Volterra model (Figure 2B, Supplemental File 1 equations 1-6). Our second model instead assumes that episymbionts compete with hosts for resources and inhibit host growth (at inhibition coefficient α) without directly killing them (Figure 2B, Supplemental File 1 equations 1-2,7-10). Both models produced similar simulated infection cycles, showing that changes to the α parameter drastically altered host crash rate, surviving host population, and stable-symbiosis density. Models also predicted that sufficiently high α values could cause long-term oscillations in host density that gradually dampen in amplitude. Since it is unclear if episymbionts kill their hosts, inhibit host growth, or some combination of the two, we consider both models in our studies.

During our experiments, we observed two rare outcomes that could happen outside a continuum ODE model: **jackpot recovery** and **total extinction** (Figure 2C). Jackpot recovery occurs when the host population manages to adapt before completely crashing, presumably due to a mutation before or during infection. Conversely, total extinction occurs when the host population fails to adapt before continuous dilution removes them from the population completely. Each rare outcome only occurred once or twice during our studies to date (38 total runs) and as such their true frequencies cannot yet be estimated. Notably, neither rare outcome would have been detectable in experiments using discontinuous, serial passaging cultivation.

Experimentally, *Nanosynbacteraceae* species’ infection cycles differed significantly, revealing a continuum of host-inhibition severity and the potential for oscillating population booms and busts (Figure 3A-B). *N. lyticus* (shown in red) and BB004 (green) both followed the standard infection cycle depicted in Figure 2A. Conversely, TM7-008 infected populations (purple) crashed and recovered two or more times with amplitude dampening between each subsequent crash; exactly as our ODE models predicted would occur for strains with a high α value. *N. lyticus* induced host population crash ∼30.0 hours post-infection (hpi), TM7-008 induced its first host crash ∼17.1 hpi, and BB004 induced host crash ∼59.2 hpi (Figure 3C). This pattern perfectly matches the pattern of competitive fitness previously observed in these strains (23), suggesting that the rate at which an episymbiont can establish host association/inhibition may dictate their competitive potential.

**Figure 3.**
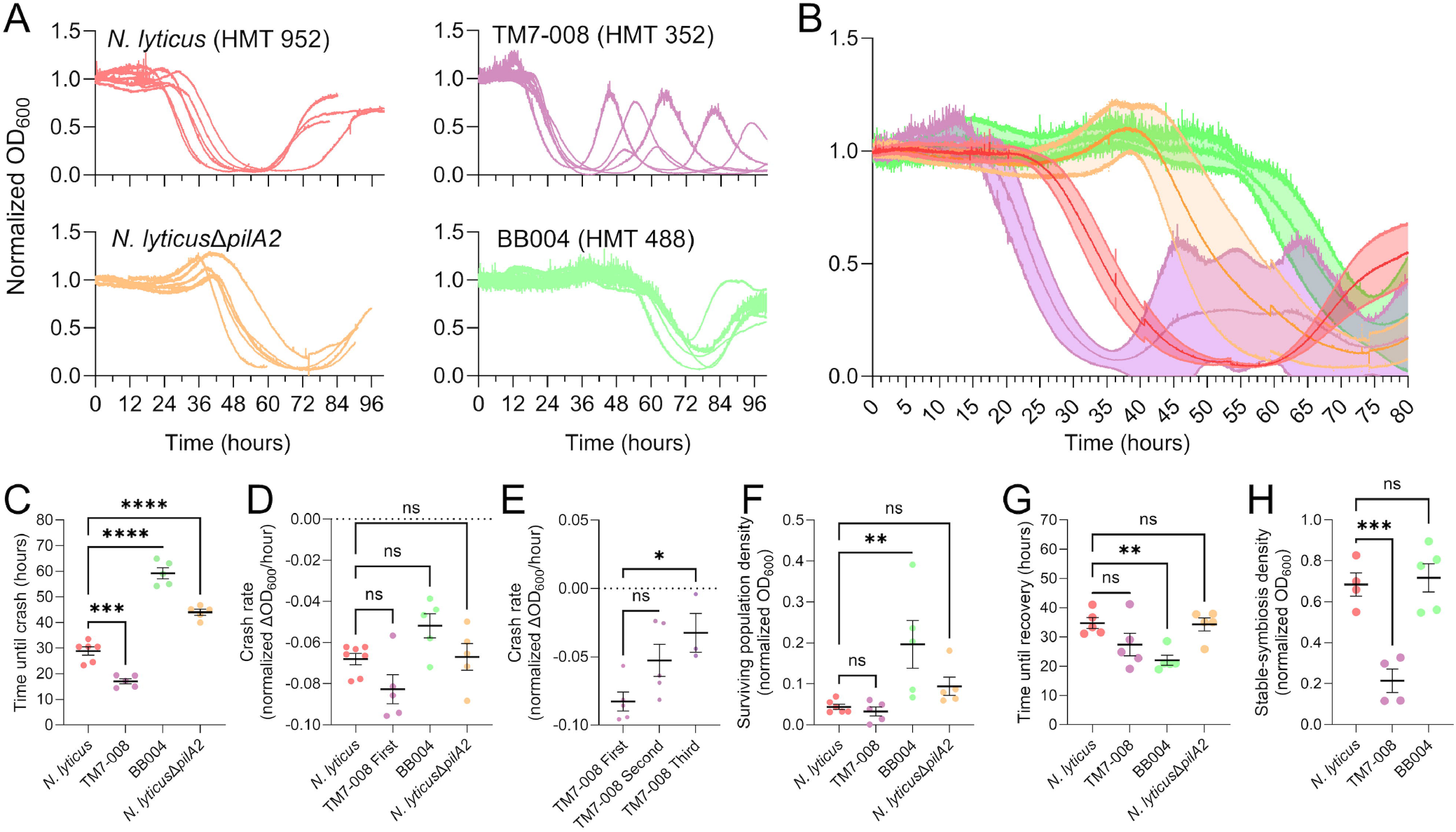
Population dynamics of constantly growing *S. odontolytica* infected by diverse episymbionts. **(A)** continuous cultivation of co-cultures infected by N. lyticus (red), *N. lyticus*Δ*pilA2* (yellow), TM7-008 (purple), and BB004 (green) demonstrate distinct infection cycles and host inhibition intensities, most clearly visible when **(B)** averaging 5-6 replicates. Most strains induce host inhibition, cause a population collapse, become subject to wash out, and then recover to stable, however TM7-008 induces two or more sequential crashes/recoveries before the host can adapt. To quantify episymbiotic impact on host populations, curves were analyzed for **(C)** time until crash, **(D)** maximum crash rate (comparing species), **(E)** maximum crash rate (comparing sequential crashes), **(F)** lowest surviving population density achieved during crash, **(G)** time until recovery, and **(H)** stable symbiosis density after recovery. The time until crash metric showed the most sensitivity and consistency between replicates.

Observed crash rates had wide intra-treatment variance and, despite having trends that resembled time until crash, did not significantly differ between species (Figure 3D, Supplemental Figure 1C-E, Supplemental Figure 2A). However, during infection by TM7-008, the “later” crashes had a significantly decreased crash rate relative to the initial host crash (Figure 3E), reflecting the gradual amplitude dampening. During the crash phase, populations infected by *N. lyticus* and TM7-008 had surviving population densities between 3-4% on average, while populations infected by BB004 had a larger average surviving population of 19.7% (Figure 3F, Supplemental Figure 2B). Furthermore, BB004 infected populations had a significantly shorter time until recovery (22.0 hours post-crash) than those infected by *N. lyticus* (34.8 hpc) (Figure 3G, Supplemental Figure 2C). These data suggest that BB004 is either poorly adapted to parasitize *S. odontolytica* XH001, or it has adapted to be less parasitic/more commensal than the other assayed species. Like crash rate, recovery rate was approximated by linear regression; however, all strains showed similar recovery rates (0.04-0.06 normalized OD_600_/hour), suggesting it is intrinsic to the basal growth rate of *S. odontolytica* and the dilution factor of the chemostat (Supplemental Figure 1G). After population recovery, *N. lyticus* and BB004 co-cultures reached similar stable-symbiosis densities at 67-68% of the host steady-state (Figure 3H). Due to constant oscillation most TM7-008 co-cultures did not reach a true stable-symbiosis density before the experiment ended, so stable-symbiosis densities were estimated by averaging their last 600 readings (50 min.). After two or three crash/recovery oscillations TM7-008 co-cultures approached a significantly lower stable-symbiosis density, at ∼21%, suggesting that TM7-008 may be more parasitic/less commensal than the other assayed species. This experimentally observed shift in stable-symbiosis density favors the host-killing ODE model because the competition model predicts that stable-symbiosis density would only vary for strains with low α values (Supplemental Figure 2D).

To better understand how a single mechanistic change can impact the episymbiont infection cycle, we utilized a *N. lyticus* mutant that is defective in long range host binding because it cannot produce one of the two type IV pili produced by *N. lyticus*. In previous studies (23), passage-based cultivation of *N. lyticus*Δ*pilA2* demonstrated reduced growth and host binding in early passages that with additional passages could eventually reach high concentrations like wildtype. We hypothesized that this strain would have an elongated time until crash but would otherwise be similar to wildtype. Within our bioreactors, *N. lyticus*Δ*pilA2* did induce a significant delay in time until crash (45.0 hpi) relative to wildtype (30.0 hpi) (Figure 3A-2C), and despite this 50% delay in establishment, the crash rate and time until recovery were unaffected (Figure 3D-G). This establishes that the time until crash metric is influenced by how long it takes an episymbiont to find and bind its host, while the other infection cycle dynamics appear to be independent phenomena. Unfortunately, stable-symbiosis density values were not determined for *N. lyticus*Δ*pilA2* replicates due to the significant crash delay.

Both of our ODE models predict that, for large values of the α parameter, the host population crashes to a small fraction of host steady-state, then undergoes decaying oscillation to the stable-symbiosis density (Figure 2B). However, the wildtype *N. lyticus* infection cycle shows a different dynamic, in which the host population crashes to a small community of survivors, then nearly recovers to the host steady-state without oscillation (Figure 3A). We hypothesized that a model that incorporated host adaptation could explain this dynamic. If a population of parasite-resistant hosts are selected for, then the population’s α parameter value decreases post-crash. We simulated this *in silico* using a modified version of the host-killing ODE model that introduced a probability (p) that hosts would become resistant post-crash (see Supplemental Files 1 and 3 for model details). In this two-host (“resistant” and “susceptible”) model, the rebound population tends to stabilize without oscillation (Figure 4A). When varying the probability of host resistance, the two-host ODE model predicts that stable-symbiosis density correlates to the percentage of resistant hosts, and that oscillation only occurs when an episymbiont’s α parameter is high and few resistant hosts emerge after the crash phase. (Figure 4B).

**Figure 4.**
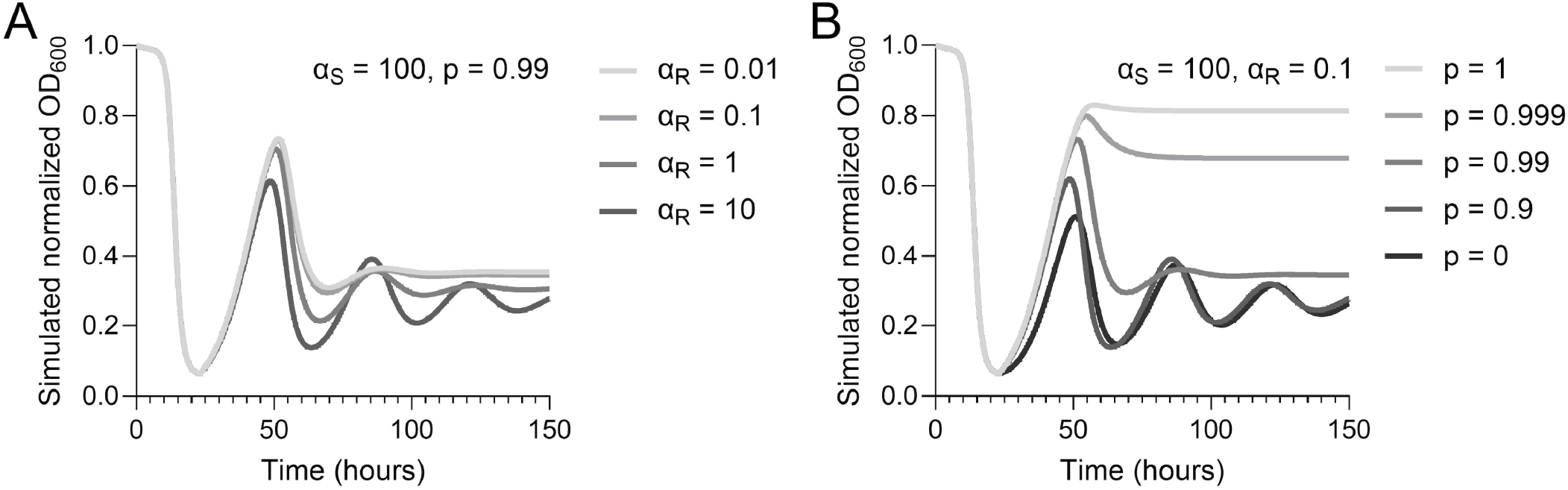
A two-host ODE model simulating selection for episymbiont resistance. Like in the host-killing episymbiont model, episymbiont susceptible hosts are initially inhibited at rate α_S_, however, after the crash susceptible hosts become resistant at rate p. Resistant hosts are not immune to episymbiont inhibition but are inhibited at the lower rate α_R_. **(A)** Simulating infection cycles while varying the α_R_ parameter shows that oscillations do not occur for sufficiently resistant host cells. **(B)** Holding α_S_ and α_R_ steady and varying the p parameter predicts that the percentage of hosts that acquire resistance strongly influences stable-symbiosis density. Furthermore, population oscillations are not predicted to occur when susceptible hosts are absent (p = 1) or rare (p > 0.99).

## Discussion

Studying episymbiont-host interactions intrinsically poses many technical challenges. These studies have focused on a single host organism and a small panel of episymbionts to develop new techniques and to quantify the ecological impact of different species and genotypes. Using this new toolset, we observed that Saccharibacteria can exist across the commensal-to-parasite continuum and have distinct impacts on host communities, but the potential for continuous cultivation to elucidate the microbial ecology of episymbiosis does not stop here. Future experiments could utilize mock communities to quantify realistic episymbiont competitions, perform continuous evolutionary experiments under changing conditions, or screen other *N. lyticus* mutants to better understand the physiological features that shape the host-inhibition factor α. A major limitation of our current experimental set up is that our Pioreactor systems do not have means of quantifying the episymbiont and host populations simultaneously. Future experiments could implement episymbiont tracking directly, by incorporating an on-bioreactor fluorometer and using fluorescent protein expressing episymbionts, or indirectly, by incorporating a fluidic autosampler and using downstream qPCR analysis to quantify episymbionts. Having real-time tracking of both populations would significantly improve our ability to model host-episymbiont dynamics. The ODE models described here are helpful for translating how episymbionts with different inhibitory potentials affect their host population; however, no model perfectly captured the dynamics of all strains. Realistically, far more data would need to be collected to determine the modes of inhibition utilized by each episymbiotic species and model how those shape each infection cycle. These models can be refined, rethought, or even combined as infection cycles from more host-episymbiont pairings are quantified. Simply by providing a continuous supply of new hosts, a more realistic environment can be recreated, and episymbiont behaviors can be made experimentally tractable in ways they never have been before.

## Methods

### Bacteria cultivation conditions

*S. odontolytica* XH001, *N. lyticus* TM7x, *N. lyticus* NS1::sfGFP (24), TM7-008, and BB004 strains were cultivated in Brain Heart Infusion (BHI) broth (Fisher: 237200) at 37 °C under microaerophilic conditions (5% CO2, 2% O2, 93% N2) in either a Whitley A35 or Coy chamber. Prior to chemostat cultivation, *N. lyticus* cells were isolated as previously described using 0.45 micron filters (9). All episymbionts were isolated from existing *S. odontolytica* containing co-cultures (13, 23). Bacterial cultures were stored at -80 °C in 15% glycerol cryo-protectant.

### Chemostat operation

All experiments were performed using four Pioreactor 20ml chemostats (hardware v1.0 and v1.5; software v25.11.19) (Pioreactor Inc. Waterloo, Canada) (11, 12). Prior to chemostat operation the reactor vial, tubing, media bottles, and waste receptacles were cleaned by cycling through 50 ml of 2% bleach and 50 ml of ultrapure water. Bleaching was essential for these experiments because minute *S. odontolytica* biofilms within the tubing can survive autoclave sterilization alone. 700 ml of BHI medium was added, then materials were autoclaved for sterility and assembled in a biosafety hood. 300 μl of overnight *S. odontolytica* culture was sterilely inoculated into the reactor via syringe and BHI was added to reach the vial’s working volume of 15 ml. 380 μl of fresh medium was pumped into the chamber every 5 min. to maintain a 0.30 dilution rate. Host cells were grown continuously for 24-36 hours to reach a steady-state OD_600_ and growth rate. Upon reaching stability, 200 μl of isolated TM7x at OD_600_ 0.005 (≈4*10^7^ cells) was injected into the reactor (roughly equivalent to an MOI 0f 0.01). Pioreactors were programmed to maintain a temperature of 37°C, spin the culture at ∼500 rpm, and take OD_600_ measurements every 5 seconds. Continuous cultures were maintained until cultures completely collapsed and, where possible, allowed to continue further into the recovery phase. Each infection treatment was replicated 5-6 times. Data were normalized to host steady-state OD_600_ to account for discrepancy between Pioreactor sensors. All data were analyzed in GraphPad Prism 10 (Version 10.3.0) via One-way ANOVA with Dunnet’s comparison. Crash rate and recovery rate were calculated via linear regression of the 6-hour period of maximum decrease/increase. A detailed step-by-step version of this protocol is available on protocols.io at https://dx.doi.org/10.17504/protocols.io.14egn5r9yg5d/v1

## Computational modelling

All details and equations for the host-killing and nutritional-competition ODE models are included in Supplemental File 1. Simulations were calculated in Python version 3.14. Code is available at https://github.com/grossmanmicrobio/Continuous-Cultivation-EpiModels or as Supplemental Files 2 and 3.

## Supporting information

Supplemental file 1 - Model Descriptions

Supplemental file 2 - Host-Killing and Competition ODE models

Supplemental file 3 - Two-host ODE Model

## Author contributions statement

Described studies were conceptualized by ASG and BB. Methodology was developed by ASG, JW, and ADS. Software was developed and run by ADS. Validation and formal analysis were performed by ASG and ADS. Investigation by ASG and JW. Data curation by ASG and BB. The original draft was written by ASG and JW. All authors contributed to review and editing. Data visualization by ASG. BB provided project administration, supervision, and funding acquisition.

## Funding and Acknowledgements

This research was supported by grants from the National Institute of Dental and Craniofacial Research (NIDCR) under Awards 1R01DE031274 (BB), 1R01DE023810 (BB), T90 DE026110-

07 (AG), and R25DE034582 (JW). Additionally, we extend our sincerest thanks to Cameron Davidson-Pilon for help troubleshooting Pioreactor assembly and operation.

---

**Supplemental figure 1.**
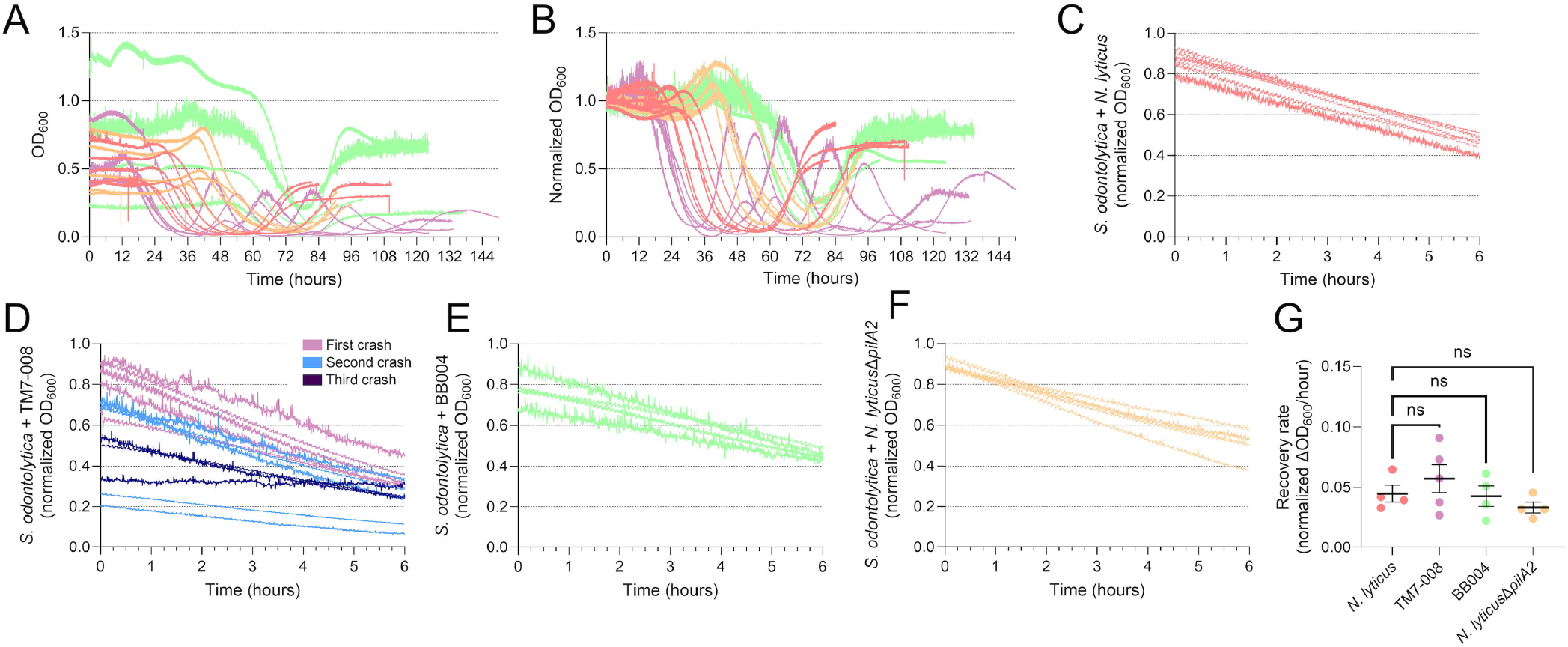
OD_600_ normalization and crash/recovery phase dynamics. **(A)** Raw OD_600_ values for all *S. odontolytica*/episymbiont co-cultures showed significant variability due to hardware variation between bioreactor models (v1.0 and v1.5 utilize different sensors). **(B)** Prior to analysis these values were normalized to host’s steady-state density for that replicate. *S. odontolytica* crash periods from co-cultures infected with **(C)** *N. lyticus*, **(D)** TM7-008, **(E)** BB004, and **(F)** *N. lyticus*Δ*pilA2* were isolated and fit to linear regressions to calculate crash rates. **(G)** Using the same linear regression analysis to calculate recovery rates showed exceptionally similar recovery rates between treatments. This similarity suggests that logarithmic recovery rate in our bioreactors is independent of episymbiont identity.

**Supplemental figure 2.**
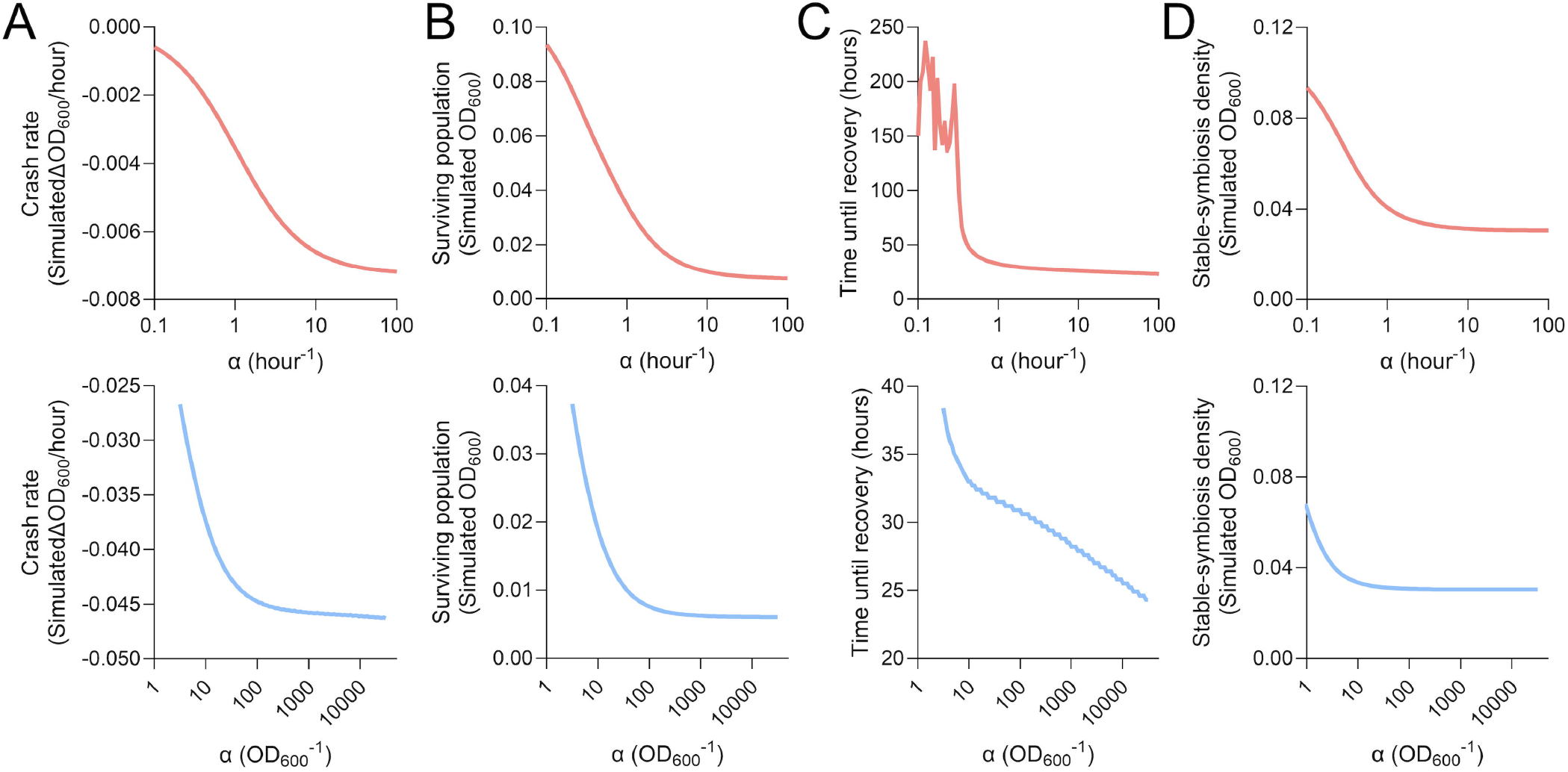
ODE simulated host-episymbiont interaction metrics. To compare the observed episymbiont infection metrics to the predictions of the host-killing ODE model (red; top row) and the competition ODE model (blue; bottom row) each metric was simulated across a range of α values. **(A)** Both ODE models predict that host crash rate decreases (i.e. becomes more negative) as α increases, before eventually approaching a plateau. **(B)** Similarly, both ODE models predict that the host population that survives the crash phase inversely correlates with α before plateauing. **(C)** When quantifying time until recovery, the host-killing ODE model predicts an almost step wise transition, with long recoveries for low α values and a rapid transition to the lower plateau as α increases. Conversely, the competition ODE model predicts that time until recovery decreases in a logarithmic, and then semi-logarithmic fashion as α increases. **(D)** Models were most strongly distinguished by their predictions for steady-state co-culture density. The host-killing model predicts that stable-symbiosis density inversely correlates with α over a wide range, while the competition model predicts that steady state density is invariant to α for all but the smallest α values.

