## Supplemental file 1 - Model Descriptions for "Resolving Host-Episymbiont Interaction Dynamics through Continuous Cultivation"

*Model definition*:

Starting with the standard cell balance over the chemostat of volume V, where fresh media is flowed in at a flowrate F. Define cell concentrations of host and episymbiont as X_H_ and X_E_, respectively, and their individual exponential growth rates as µ_H_ and µ_E_ and carrying capacities K_H_ and K_E_.

The uncoupled logistic growth equations for each species are (1a) and (2a)

$V\frac{dX_{H}}{dt}=\mu_{H}X_{H}V\left( 1-\frac{X_{H}}{K_{H}} \right)-FX_{H}$	(1a)

$V\frac{dX_{E}}{dt}=\mu_{E}X_{E}V\left( 1-\frac{X_{E}}{K_{E}} \right)-FX_{E}$	(2a)

We define the dilution rate D as the ratio between the flow rate and volume and simplify these ODEs as:

$\frac{dX_{H}}{dt}=\mu_{H}X_{H}\left( 1-\frac{X_{H}}{K_{H}} \right)-{DX}_{H}$	(1b)

$\frac{dX_{E}}{dt}=\mu_{E}X_{E}\left( 1-\frac{X_{E}}{K_{E}} \right)-DX_{E}$	(2b)

We initially considered two models for the host-episymbiont coupling on growth rate. In the first model (“Killing Model”), the episymbiont actively kills the host but its own growth rate is proportional to the host population, similar to a Lotka-Volterra model. In this case, the coupled differential equation model is

$\frac{dX_{H}}{dt}=\mu_{H}X_{H}\left( 1-\frac{X_{H}}{K_{H}} \right)-DX_{H}-\alpha X_{E}X_{H}$	(3)

$\frac{dX_{E}}{dt}=\frac{\mu_{E}X_{E}X_{H}}{K_{H}}\left( 1-\frac{X_{E}}{K_{E}} \right)-DX_{E}$	(4)

where the new parameter α is a second-order rate constant for host cell death.

A single positive steady-state cell concentration for both species can be directly calculated from this model by setting the differential terms to zero and solving the algebraic system of equations:

$X_{H,SS}=K_{H}\frac{K_{E}\alpha\mu_{E}+\mu_{E}D-\mu_{E}\mu_{H}+\sqrt{\left( K_{E}\alpha\mu_{E}+\mu_{E}D-\mu_{E}\mu_{H} \right)^{2}-4K_{E}\alpha\mu_{E}\mu_{H}D}}{2\mu_{H}\mu_{E}}$	(5)

$X_{E,SS}=K_{E}(1-\frac{K_{H}D}{\mu_{E}X_{H,SS}})$	(6)

It is easy to see here that the host steady state declines as alpha increases, and this decline also reduces the episymbiont steady-state concentration, relative to each species’ carrying capacity.

In the second model (“Competition Model”), the episymbiont outcompetes the host for resources and inhibits its growth without directly killing it. This generates the model as follows:

$\frac{dX_{H}}{dt}=\frac{\mu_{H}X_{H}\left( 1-\frac{X_{H}}{K_{H}} \right)}{1+\alpha X_{E}}-DX_{H}$	(7)

$\frac{dX_{E}}{dt}=\frac{\mu_{E}X_{E}X_{H}}{K_{H}}\left( 1-\frac{X_{E}}{K_{E}} \right)-DX_{E}$	(8)

Now, the α term is analogous to the inhibition coefficient for an enzyme and has units of inverse-cells. The greater the value of alpha, the lower the concentration of episymbiont is required to inhibit the growth of the host. Again, at steady state, this quadratic equation can be solved for the positive roots, yielding:

$X_{E,SS}=\frac{\mu_{E}\mu_{H}-D+\alpha DK_{E}\mu_{E}-\sqrt{\left( \mu_{E}\mu_{H}-D+\alpha DK_{E}\mu_{E} \right)^{2}+4\alpha\mu_{E}D(\mu_{H}DK_{E}+DK_{E}\mu_{E}-\mu_{E}\mu_{H}K_{E})}}{2\alpha\mu_{E}D}$	(9)

$X_{H,SS}=\frac{K_{H}}{\mu_{H}}(\mu_{H}-D-\alpha X_{E,SS}D)$	(10)

Conversely, in Model 2, the episymbiont steady-state population decreases as alpha increases. However, this does not necessarily lead to either an increase or a decrease in the host steady-state population.

Based on prior experiments, we choose the following parameter set and vary α.

For Supplemental Figure 3, we considered a simple two-host predation model where the host gains resistance to the parasite during the initial crash phase, which leads to an effective decrease in alpha, the killing rate of the parasite (but it is still nonzero).

Consider the predation model but one where the host gains resistance to the parasite during the initial crash phase, which leads to an effective decrease in alpha, the killing rate of the parasite (but it is still nonzero.) In this case, we treat alpha in the coupled model as

$\frac{dX_{H}}{dt}=\mu_{H}X_{H}\left( 1-\frac{X_{H}}{K_{H}} \right)-DX_{H}-\alpha X_{E}X_{H}$	(3)

$\frac{dX_{E}}{dt}=\frac{\mu_{E}X_{E}X_{H}}{K_{H}}\left( 1-\frac{X_{E}}{K_{E}} \right)-DX_{E}$	(4)

$\alpha= \left\{ \begin{matrix} \alpha_{S} & if t<t_{crash} \\ p\alpha_{S}+\left( 1-p \right)\alpha_{R} & if t>t_{crash} \end{matrix} \right\}$	(11)

where *p* represents the probability of a mutant emerging.
