## Supplemental file 2 - Host-Killing and Competition ODE models for "Resolving Host-Episymbiont Interaction Dynamics through Continuous Cultivation": SupplementalFile2_piorxtr_ODE.ipynb

```
# %% [code]
import numpy as np
import seaborn as sns
import matplotlib.pyplot as plt
import math
import scipy as spy
import pandas as pd

# %% [markdown]
Let's start with a model where the symbiont actively kills the host.


$$\frac{dX_H}{dt}=\mu_H X_H (1 - \frac{X_H}{K_H}) - DX_H - \alpha X_H X_H$$
$$\frac{dX_E}{dt}=\mu_E X_H X_E (1- \frac{X_E}{K_E}) -D X_E $$
This more resembles a Lotka-Volterra type dynamic.

# %% [code]
def dX_modelkilling(X,t,K_1, K_2, k, mu_1,mu_2,D,alpha):
    X1=X[0]
    X2=X[1]
    dX1 = mu_1 * X1 * (1 - X1/K_1) - D*X1 - alpha*X1*X2
    dX2 =  mu_2 * X1 * X2 * (1 - X2/K_2) - D * X2
    return [dX1,dX2]

K_1 = 0.356
D = 0.304
mu_1 = 0.445
K_2 = 0.356
t_start = 0
t_end = 150
mu_2 = 10


t_fit = np.linspace(t_start,t_end,10000)
alpha_list = [0,0.01,0.1,1,10,100]

write_arr = np.zeros((len(t_fit),2*len(alpha_list)+1))
write_arr[:,0] = t_fit
key_list = ['time (hrs)']

for i in range(len(alpha_list)):
    X = spy.integrate.odeint(dX_modelkilling,[0.115,1E-7],t_fit,args=(K_1,K_2,k,mu_1,mu_2,D,alpha_list[i]))
    plt.plot(t_fit,(X[:,0]),label=r'$\alpha$ = {0}'.format(alpha_list[i]))
    write_arr[:,2*i+1] = X[:,0]
    write_arr[:,2*i+2] = X[:,1]
    key_list.append('Xh, alpha = ' + str(alpha_list[i]))
    key_list.append('Xe, alpha = ' + str(alpha_list[i]))
plt.xlabel('Time (hrs)')
plt.ylabel('Normalized OD of Host')
plt.legend()
plt.title('Host OD vs. Predation Rate, Killing Model')
plt.savefig('Host OD vs. Alpha, Killing Model.jpg')
df_write = pd.DataFrame(data=write_arr,columns=key_list)
df_write.to_excel('Killing Model Trajectories.xlsx')
#plt.clf()


# %% [code]
def dX_model_competitive(X,t,K_1, K_2, k, mu_1,mu_2,D,alpha):
    X1=X[0]
    X2=X[1]
    dX1 = mu_1 * X1 * (1 - X1/K_1) / (1+alpha*X2) - D*X1
    dX2 =  mu_2 * X1 * X2 * (1 - X2/K_2) - D * X2
    return [dX1,dX2]

K_1 = 0.356
D = 0.304
mu_1 = 0.445
K_2 = 0.356
t_start = 0
t_end = 150
mu_2 = 10


t_fit = np.linspace(t_start,t_end,10000)
alpha_list = [0,0.01,0.1,1,10,100,1000]

write_arr = np.zeros((len(t_fit),2*len(alpha_list)+1))
write_arr[:,0] = t_fit
key_list = ['time (hrs)']

for i in range(len(alpha_list)):
    X = spy.integrate.odeint(dX_model_competitive,[0.115,1E-7],t_fit,args=(K_1,K_2,k,mu_1,mu_2,D,alpha_list[i]))
    plt.plot(t_fit,(X[:,0]),label=r'$\alpha$ = {0}'.format(alpha_list[i]))
    write_arr[:,2*i+1] = X[:,0]
    write_arr[:,2*i+2] = X[:,1]
    key_list.append('Xh, alpha = ' + str(alpha_list[i]))
    key_list.append('Xe, alpha = ' + str(alpha_list[i]))
plt.xlabel('Time (hrs)')
plt.ylabel('Normalized OD of Host')
plt.legend()
plt.title('Host OD vs. Inhibition Factor Rate, Competition Model')
plt.savefig('Host OD vs. Alpha, Competition Model.jpg')
df_write = pd.DataFrame(data=write_arr,columns=key_list)
df_write.to_excel('Competition Model Trajectories.xlsx')
#plt.clf()


# %% [code]
#instead of plotting trajectories, let's plot derivatives (using a five point stencil):

def dX_modelkilling(X,t,K_1, K_2, k, mu_1,mu_2,D,alpha):
    X1=X[0]
    X2=X[1]
    dX1 = mu_1 * X1 * (1 - X1/K_1) - D*X1 - alpha*X1*X2
    dX2 =  mu_2 * X1 * X2 * (1 - X2/K_2) - D * X2
    return [dX1,dX2]

def fivepointstencil(x):
    return [x[i]-8*x[i+1]+8*x[i+3]-x[i+4] for i in range(len(x)-5)]
    

K_1 = 0.356
D = 0.304
mu_1 = 0.445
K_2 = 0.356
t_start = 0
t_end = 150
mu_2 = 10


t_fit = np.linspace(t_start,t_end,10000)
alpha_list = [0,0.01,0.1,1,10,100]


for i in range(len(alpha_list)):
    X = spy.integrate.odeint(dX_modelkilling,[0.115,1E-7],t_fit,args=(K_1,K_2,k,mu_1,mu_2,D,alpha_list[i]))
    #plt.plot(t_fit,(X[:,0]),label=r'$\alpha$ = {0}'.format(alpha_list[i]))
    plt.plot(t_fit[3:-2],fivepointstencil(X[:,0]),label=r'$\alpha$ = {0}'.format(alpha_list[i]))


plt.xlabel('Time (hrs)')
plt.ylabel('Time Rate of Change of Host Concentration')
plt.legend()
plt.title('dX/dt vs. Predation Rate, Killing Model')

#plt.clf()


# %% [code]
#thresholds: time derivative has to exceed -0.0001 to count as a crash. Crash rate is the slope at the inflection point

def dX_modelkilling(X,t,K_1, K_2, k, mu_1,mu_2,D,alpha):
    X1=X[0]
    X2=X[1]
    dX1 = mu_1 * X1 * (1 - X1/K_1) - D*X1 - alpha*X1*X2
    dX2 =  mu_2 * X1 * X2 * (1 - X2/K_2) - D * X2
    return [dX1,dX2]

def fivepointstencil(x):
    return np.array([x[i]-8*x[i+1]+8*x[i+3]-x[i+4] for i in range(len(x)-5)])

K_1 = 0.356
D = 0.304
mu_1 = 0.445
K_2 = 0.356
t_start = 0
t_end = 300
mu_2 = 10

threshold = -0.0001


t_fit = np.linspace(t_start,t_end,10000)
alpha_list = np.logspace(-1,2,100) #alpha must exceed 0.1 or there is no crash

all_data = np.zeros((len(alpha_list),6))
all_data[:,0]=alpha_list


key_list = ['alpha','time to crash (hrs)','crash rate (hr^-1)','surviving population (OD)','time to recovery (hrs)','steady state population (OD)']

for i in range(len(alpha_list)):
    X = spy.integrate.odeint(dX_modelkilling,[0.115,1E-7],t_fit,args=(K_1,K_2,k,mu_1,mu_2,D,alpha_list[i]))
    dX = fivepointstencil(X[:,0])
    t_index = t_fit[3:-2]
    time_to_crash = t_index[dX<threshold][0]
    min_OD = np.min(X[:,0])
    time_min_OD = t_fit[np.argmin(X[:,0])]
    crash_rate = np.min(dX)
    time_of_recovery = t_index[np.logical_and(t_index>time_min_OD,dX>0)][0]
    time_till_recovery = time_of_recovery-time_to_crash
    steady_state = np.mean(X[-50:,0])

    all_data[i,1]=time_to_crash
    all_data[i,2] = crash_rate
    all_data[i,3] = min_OD
    all_data[i,4] = time_till_recovery
    all_data[i,5] = steady_state
    

df_write = pd.DataFrame(data=all_data,columns=key_list)
df_write.to_excel('Killing Model Summary Data.xlsx')

for i in range(5):
    fig= plt.subplot(2,3,i+1)
    plt.plot(alpha_list,all_data[:,i+1],marker='.')
    plt.xlabel('alpha',fontsize='small')
    plt.xscale('log')
    plt.ylabel(key_list[i+1],fontsize='small')
    plt.title(str(key_list[i+1]),fontsize='small')
plt.tight_layout()
plt.savefig('Killing Model Summary Data.jpg')


# %% [code]
#thresholds: time derivative has to exceed -0.0001 to count as a crash. Crash rate is the slope at the inflection point

def dX_model_competitive(X,t,K_1, K_2, k, mu_1,mu_2,D,alpha):
    X1=X[0]
    X2=X[1]
    dX1 = mu_1 * X1 * (1 - X1/K_1) / (1+alpha*X2) - D*X1
    dX2 =  mu_2 * X1 * X2 * (1 - X2/K_2) - D * X2
    return [dX1,dX2]

def fivepointstencil(x):
    return np.array([x[i]-8*x[i+1]+8*x[i+3]-x[i+4] for i in range(len(x)-5)])

K_1 = 0.356
D = 0.304
mu_1 = 0.445
K_2 = 0.356
t_start = 0
t_end = 300
mu_2 = 10

threshold = -0.0001


t_fit = np.linspace(t_start,t_end,10000)
alpha_list = np.logspace(0.5,4.5,100) #alpha must exceed 0.5 or there is no crash

all_data = np.zeros((len(alpha_list),6))
all_data[:,0]=alpha_list


key_list = ['alpha','time to crash (hrs)','crash rate (hr^-1)','surviving population (OD)','time to recovery (hrs)','steady state population (OD)']

for i in range(len(alpha_list)):
    X = spy.integrate.odeint(dX_model_competitive,[0.115,1E-7],t_fit,args=(K_1,K_2,k,mu_1,mu_2,D,alpha_list[i]))
    dX = fivepointstencil(X[:,0])
    t_index = t_fit[3:-2]
    time_to_crash = t_index[dX<threshold][0]
    min_OD = np.min(X[:,0])
    time_min_OD = t_fit[np.argmin(X[:,0])]
    crash_rate = np.min(dX)
    time_of_recovery = t_index[np.logical_and(t_index>time_min_OD,dX>0)][0]
    time_till_recovery = time_of_recovery-time_to_crash
    steady_state = np.mean(X[-50:,0])

    all_data[i,1]=time_to_crash
    all_data[i,2] = crash_rate
    all_data[i,3] = min_OD
    all_data[i,4] = time_till_recovery
    all_data[i,5] = steady_state
    

df_write = pd.DataFrame(data=all_data,columns=key_list)
df_write.to_excel('Competitive Model Summary Data.xlsx')

for i in range(5):
    fig= plt.subplot(2,3,i+1)
    plt.plot(alpha_list,all_data[:,i+1],marker='.')
    plt.xlabel('alpha',fontsize='small')
    plt.xscale('log')
    plt.ylabel(key_list[i+1],fontsize='small')
    plt.title(str(key_list[i+1]),fontsize='small')
plt.tight_layout()
plt.savefig('Competitive Model Summary Data.jpg')


# %% [code]
```
