## Supplemental file 3 - Two-host ODE Model for "Resolving Host-Episymbiont Interaction Dynamics through Continuous Cultivation": SupplementalFile3_dynamic_alpha.ipynb

```
# %% [markdown]
Consider the predation model but one where the host gains resistance to the parasite during the initial crash phase, which leads to an effective decrease in alpha, the killing rate of the parasite (but it is still nonzero.)
$$\frac{dX_H}{dt}=\mu_H X_H (1 - \frac{X_H}{K_H}) - DX_H - \alpha X_E X_H$$
$$\frac{dX_E}{dt}=\mu_E X_H X_E (1- \frac{X_E}{K_E}) -D X_E $$

where 

$\alpha = \begin{cases}
\alpha_H & \text{if } t < t_{crash},\\
p\alpha_L + (1-p)\alpha_H & \text{if } t > t_{crash}\\
\end{cases}$

where $p$ represents the probability of a mutant emerging.


# %% [code]
import numpy as np
import seaborn as sns
import matplotlib.pyplot as plt
import math
import scipy as spy
import pandas as pd

def dX_dynamic_killing(X,t,K_1, K_2, k, mu_1,mu_2,D,alpha_H,alpha_L,p,t_crash):
    X1=X[0]
    X2=X[1]
    if t>t_crash:
        alpha = p*alpha_L + (1-p)*alpha_H
    else:
        alpha = alpha_H
    dX1 = mu_1 * X1 * (1 - X1/K_1) - D*X1 - alpha*X1*X2
    dX2 =  mu_2 * X1 * X2 * (1 - X2/K_2) - D * X2
    return [dX1,dX2]


def dX_static_killing(X,t,K_1, K_2, k, mu_1,mu_2,D,alpha):
    X1=X[0]
    X2=X[1]
    dX1 = mu_1 * X1 * (1 - X1/K_1) - D*X1 - alpha*X1*X2
    dX2 =  mu_2 * X1 * X2 * (1 - X2/K_2) - D * X2
    return [dX1,dX2]

def fivepointstencil(x):
    return np.array([x[i]-8*x[i+1]+8*x[i+3]-x[i+4] for i in range(len(x)-5)])

K_1 = 0.356
D = 0.304
mu_1 = 0.445
K_2 = 0.356
t_start = 0
t_end = 150
mu_2 = 10
alphaH_list = [100,100,100,100]
alphaL_list = [0.01,0.1,1,10]
p_list = [0.99,0.99,0.99,0.99]
k=1


t_fit = np.linspace(t_start,t_end,10000)
write_arr = np.zeros((len(t_fit),2*len(p_list)+1))
write_arr[:,0] = t_fit
key_list = ['time (hrs)']

for i in range(len(alphaH_list)):
    #find time to crash with alpha not changing
    X_static = spy.integrate.odeint(dX_static_killing,[0.115,1E-7],t_fit,args=(K_1,K_2,k,mu_1,mu_2,D,alphaH_list[i]))
    dX_static = fivepointstencil(X_static[:,0])
    t_index = t_fit[3:-2]
    crash_time = t_fit[np.argmin(X_static[:,0])]

    #now rerun simulation with this crash time incorporated
    X_dynamic = spy.integrate.odeint(dX_dynamic_killing,[0.115,1E-7],t_fit,args=(K_1,K_2,k,mu_1,mu_2,D,alphaH_list[i],alphaL_list[i],p_list[i],crash_time))

    plt.plot(t_fit,(X_dynamic[:,0]),label=r'$\alpha_H$ = {0},$\alpha_L$ = {1}, p = {2}'.format(alphaH_list[i],alphaL_list[i],p_list[i]))
    #plt.plot(t_fit,(X_dynamic[:,1]),label=r'PARASITE $\alpha_H$ = {0},$\alpha_L$ = {1}, p = {2}'.format(alphaH_list[i],alphaL_list[i],p_list[i]))

    write_arr[:,2*i+1] = X_dynamic[:,0]
    write_arr[:,2*i+2] = X_dynamic[:,1]
    key_list.append('host concentration, alphaL = '  + str(alphaL_list[i]))
    key_list.append('parasite concentration, alphaL = '  + str(alphaL_list[i]))
plt.xlabel('Time (hrs)')
plt.ylabel('Normalized Host OD')
plt.legend()
plt.title('Host OD vs. Predation Rates, Adaptive Killing Model')
plt.savefig('Host OD vs. Resistance, Adaptive Killing Model.jpg')
df_write = pd.DataFrame(data=write_arr,columns=key_list)
df_write.to_excel('Dynamic Killing Model Trajectories_vary_alphaL.xlsx')
#plt.clf()


# %% [code]
import numpy as np
import seaborn as sns
import matplotlib.pyplot as plt
import math
import scipy as spy
import pandas as pd

def dX_dynamic_killing(X,t,K_1, K_2, k, mu_1,mu_2,D,alpha_H,alpha_L,p,t_crash):
    #crash flag is True if the first crash has already happened
    X1=X[0]
    X2=X[1]
    if t>t_crash:
        alpha = p*alpha_L + (1-p)*alpha_H
    else:
        alpha = alpha_H
    dX1 = mu_1 * X1 * (1 - X1/K_1) - D*X1 - alpha*X1*X2
    dX2 =  mu_2 * X1 * X2 * (1 - X2/K_2) - D * X2
    return [dX1,dX2]


def dX_static_killing(X,t,K_1, K_2, k, mu_1,mu_2,D,alpha):
    X1=X[0]
    X2=X[1]
    dX1 = mu_1 * X1 * (1 - X1/K_1) - D*X1 - alpha*X1*X2
    dX2 =  mu_2 * X1 * X2 * (1 - X2/K_2) - D * X2
    return [dX1,dX2]

def fivepointstencil(x):
    return np.array([x[i]-8*x[i+1]+8*x[i+3]-x[i+4] for i in range(len(x)-5)])

K_1 = 0.356
D = 0.304
mu_1 = 0.445
K_2 = 0.356
t_start = 0
t_end = 150
mu_2 = 10
alphaH_list = [100,100,100,100,100]
alphaL_list = [0.1,0.1,0.1,0.1,0.1]
p_list = [1,0.999,0.99,0.9,0]
k=1


t_fit = np.linspace(t_start,t_end,10000)
write_arr = np.zeros((len(t_fit),2*len(p_list)+1))
write_arr[:,0] = t_fit
key_list = ['time (hrs)']

for i in range(len(alphaH_list)):
    #find time to crash with alpha not changing
    X_static = spy.integrate.odeint(dX_static_killing,[0.115,1E-7],t_fit,args=(K_1,K_2,k,mu_1,mu_2,D,alphaH_list[i]))
    dX_static = fivepointstencil(X_static[:,0])
    t_index = t_fit[3:-2]
    crash_time = t_fit[np.argmin(X_static[:,0])]
    #now rerun simulation with this crash time incorporated
    X_dynamic = spy.integrate.odeint(dX_dynamic_killing,[0.115,1E-7],t_fit,args=(K_1,K_2,k,mu_1,mu_2,D,alphaH_list[i],alphaL_list[i],p_list[i],crash_time))

    plt.plot(t_fit,(X_dynamic[:,0]),label=r'$\alpha_H$ = {0},$\alpha_L$ = {1}, p = {2}'.format(alphaH_list[i],alphaL_list[i],p_list[i]))
    #plt.plot(t_fit,(X_dynamic[:,1]),label=r'PARASITE $\alpha_H$ = {0},$\alpha_L$ = {1}, p = {2}'.format(alphaH_list[i],alphaL_list[i],p_list[i]))

    write_arr[:,2*i+1] = X_dynamic[:,0]
    write_arr[:,2*i+2] = X_dynamic[:,1]

    key_list.append('host concentration, p = '  + str(p_list[i]))
    key_list.append('parasite concentration, p = '  + str(p_list[i]))

plt.xlabel('Time (hrs)')
plt.ylabel('Normalized Host OD')
plt.legend()
plt.title('Host OD vs. Mutation Rate, Adaptive Killing Model')
plt.savefig('Host OD vs. Mutation Rate, Adaptive Killing Model.jpg')
df_write = pd.DataFrame(data=write_arr,columns=key_list)
df_write.to_excel('Dynamic Killing Model Trajectories_vary_P.xlsx')
#plt.clf()


# %% [code]
```
